# Hormonal contraceptives alter amphetamine place preference and responsivity in the intact female rat

**DOI:** 10.1101/2021.07.28.454029

**Authors:** Emily N. Hilz, Marcelle Olvera, Dohyun Jun, Megha Chadha, Ross Gillette, Marie-H. Monfils, Andrea C. Gore, Hongjoo J. Lee

**Affiliations:** Department of Psychology, The University of Texas at Austin, Austin TX USA; Institute for Neuroscience, The University of Texas at Austin, Austin TX USA.; Division of Pharmacology and Toxicology, The University of Texas at Austin, Austin TX USA

**Keywords:** hormonal contraceptives, females, estrous cycle, amphetamine, conditioned place preference, dopamine, ultrasonic vocalizations, addiction

## Abstract

Hormonal contraceptives (HCs) are commonly used among reproductive aged women and alter the physiological state of the user by interfering with endogenous hormone concentrations and their actions on the reproductive tract. As hormones such as estradiol and progesterone modulate the incidence of substance abuse disorders in women, it is important to consider the influence HCs have on the female brain and behavior. This experiment explores how female sex steroid hormonal states associated with the rat estrous cycle, and modulating those states with HCs, influences measures of drug preference and responsivity. First, rats underwent food-light Pavlovian conditioning to measure conditioned orienting, a known predictor of amphetamine (AMP) place preference. Then, rats were conditioned and tested for AMP place preference with either an HC-implant or during estrous cycle stages associated with different ovarian hormone levels (i.e., proestrus (P) or metestrus/diestrus (M/D) while recording ultrasonic vocalizations (USVs) as an index of hedonic responsivity. Because of dopamine’s (DA) role in modulation of AMP actions, DA cell activity and availability were examined using tyrosine hydroxylase and FOS immunohistochemistry after final AMP challenge. Conditioned orienting did not differ between cycling and HC-implanted. P rats emitted more USVs during conditioning, showed higher AMP place preference throughout testing, and had higher DA cell activity in the substantia nigra compared to M/D and HC-implanted rats. Sex steroid hormone serum concentration and uterine horn thickness predicted some but not all of these measures. This experiment suggests ovarian hormones affect drug preference and responsivity, while providing novel insight into how hormone-altering contraceptives may reduce these measures.

## Introduction

The use of hormonal contraceptives (HCs) is ever-increasing among women in the United States (Kavanaugh and Jerman, 2018). HCs achieve contraception by acting directly upon the reproductive tract as well as altering the manner and rate that gonadal hormones are secreted by the female body. These hormones also have effects that extend beyond the control of reproduction; however, few studies have sought to characterize the influence of HCs on the female brain and behavior. The HC levonorgestrel (LNG) is commonly used in both birth control pills and HC implants and, like many progestin contraceptives, induces negative feedback on the release of gonadotropin releasing hormone (GnRH) from the hypothalamus and interferes with the release of sex steroid hormones that are necessary for ovulation (Rivera et al., 1999).

Estradiol and progesterone have received much attention for their opposing roles in female reward- and drug-seeking behavior: estradiol enhances female drug-seeking and drug-responsivity both endogenously over the hormonal cycle and when administered exogenously (Carroll and Anker, 2010; Anker et al., 2007), while progesterone has notable attenuating effects (Quinones-Jenab and Jenab, 2010; Feltenstein and See, 2007). Both estradiol and progesterone generally enhance dopamine (DA) neurotransmission (Diekhof, 2018; Sun et al., 2016; Becker and Rudick, 1999; Mani et al., 1996; Becker, 1990). DA facilitates reproductive behavior through its interactions progesterone and progestin receptors in the brain (Mani and Oyola, 2012; Mani et al., 1994a,b) and also contributes to reward learning and the incentive-motivation for drugs of abuse in a way that is modulated by estradiol (Alikaya et al., 2018; Shultz, 2013; Berridge, 2012; Zhang et al., 2008; Robinson et al., 2005). Because of this, female sex steroid hormones have been posited to underlie a notable increased risk of substance abuse disorder development in women compared to men (Yoest, 2014; Lynch et al., 2002).

Sex steroids have also been implicated in contributing to extinction of Pavlovian behavior: reducing sex steroid availability via ovariectomy (OVX) or with LNG impairs extinction of fear learning (Graham and Daher, 2016; Hwang et al., 2015; Graham and Milad, 2013; Milad et al., 2009); estradiol treatment enhances while endogenous progesterone attenuates extinction of conditioned drug preference (Yousuf et al., 2019; Twining et al., 2013; Feltenstein and See, 2007). Resurgence of previously extinguished appetitive or drug-seeking behaviors has also been shown to be mediated by sex steroids and ovarian hormonal states (Hilz et al., 2019b; Anderson and Petrovitch, 2015; Becker and Hu, 2008), with estradiol facilitating and progesterone attenuating return of extinguished drug-seeking behavior (Doncheck et al., 2018; Anker et al., 2007). As extinction is a common therapeutic intervention for treating drug dependency, its modulation by female sex steroid hormones is an important factor to consider when designing therapeutic interventions for women with problematic drug use.

Conditioned orienting is a form of cue-directed behavior similar to sign-tracking that is thought to represent increased incentive and/or motivational processing of conditioned stimulus (CS) information (Olshavsky et al., 2014; Holland, 1977) and to rely on functional activity in the DA system (Lee et al., 2011; 2005; Han et al., 1997; Gallagher et al., 1990). Conditioned orienting predicts female drug preference (Hilz et al., 2019a); however, little is known about the effects of ovarian hormones on conditioned orienting or how these measures may interact to affect drug preference. Studies comparing drug preference between male and female animals using conditioned place preference (CPP) methodologies generally support the supposition that females exhibit increased drug preference compared to males and have shown that estradiol increases (Mirbaha et al., 2009; Silverman and Koenig, 2007; Russo et al., 2003a,b) whereas progesterone attenuates (Russo et al., 2008; Wu et al., 2008) female drug CPP; when given in conjunction, estradiol and progesterone potentiate the magnitude of female drug CPP (Russo et al., 2008a). The literature around female drug CPP has failed to address the role of endogenous sex steroid hormonal states; moreover, the use of OVX to compare high and low sex steroid hormone treatments in female rats is not physiologically analogous to reproductive aged women (other than the small subset with oophorectomy) with substance abuse disorders or undergoing treatment for such. The present study fills this gap and provides a novel method for assessing suppressed sex steroid hormone states in female drug learning.

## Methods

### Subjects

Thirty-three female Sprague-Dawley rats (Envigo), weighing ∼200-250g upon arrival were housed in pairs on a 14:10 hour reverse light-dark cycle with lights off at 10AM. Rats were vaginally lavaged daily to monitor estrous cycle activity. Water was always available. Prior to light-food conditioning rats were food-restricted for one week to reach ∼90% body weight and maintained this restriction throughout that procedure. After light-food conditioning, rats were allowed one week for weight recovery and were subsequently fed *ad libitum*. All procedures and animal handling were conducted at the beginning of the dark cycle, were approved by the Institutional Animal Care and Use Committee at the University of Texas at Austin, and were conducted in accordance with NIH guidelines.

### Apparatus

Light-food conditioning occurred in a typical conditioning chamber measuring 30.5cm W/ 25.5cm L / 30.5cm H (Coulbourn Instruments, Whitehall, PA). Briefly, the chambers were constructed of clear acrylic front and back walls, steel-rod floors, and aluminum sides and ceiling. Food-pellets (45mg TestDiet, Richmond, IN) were dispensed into a foodcup on the right wall from an external magazine, and entries into the foodcup were measured via breaks in an infrared beam at the opening. A 2-watt bulb was located 20cm above the foodcup, and illumination of this light served as the CS for conditioning. Chambers were enclosed in sound- and light-attenuating boxes (Coulbourn Instruments, Whitehall, PA). Mounted inside the boxes but outside the chambers were digital video cameras (KT&C USA, Fairfield, NJ) used to monitor orienting activity during conditioning.

Conditioned place preference procedures occurred in a conditioning chamber similar in construction but of different dimensions (50.8cm W / 25.5cm L / 29.2cm H; Coulbourn). Walls had no perforations and an aluminum wall with a retractable door bisected the chamber into separate compartments. The door was closed during amphetamine and saline conditioning, and open during baseline / preference testing / reinstatement. Black and white paper was secured to the acrylic walls of each compartment such that one side (left) was “dark” and the other (right) was “light”. A 2-watt red light located 22.5cm above the floor provided ambient light in the dark compartment. A down-facing camera was mounted atop the dark compartment which recorded time spent by the rat within each compartment. Subtracting time spent in the dark chamber from the total session length allowed determination of time spent in the white chamber and subsequent generation of place preference scores. CM16 ultrasonic microphones (Avisoft Bioacoustics, Berlin, Germany) were located outside of the dark conditioning chamber during tests and in the opposite chamber during conditioning. Boxes were enclosed in sound- and light-attenuating boxes.

### Procedure

#### Estrous tracking

All rats received daily vaginal lavage at the beginning of the dark cycle prior to procedures. The vaginal cavity was flushed with 40µl 0.9% sterile saline, and cytological samples were immediately examined under 10X magnification to determine estrous cycle stage. Stages of the estrous cycle can be determined by the structure and quantity of vaginal epithelial cells (Goldman et al., 2007). The method of determining each stage is detailed elsewhere (see Hilz et al., 2019a,b). In short, samples containing a high quantity of primarily nucleated epithelial cells were classified as proestrus (P), of primarily cornified epithelial cells as estrus (E), of a mixture of cornified and leukocytes as metestrus (M), and of primarily leukocytes as diestrus (D). For the purposes of the CPP experiment rats were conditioned during either P or M/D to capture differential hormone states. P is associated with peaks in estradiol and progesterone (and is when ovulation occurs), while M and early D are associated with comparably lower levels of estradiol and progesterone (for review see Butcher et al., 1974). E was excluded from comparisons as hormone levels fluctuate widely throughout the stage and can be difficult to predict using cytology.

#### Levonorgestrel implant surgery

After ensuring all rats had normal estrous cycling, a random selection of rats (n = 10) were implanted with subcutaneous levonorgestrel (LNG) capsules. LNG implant was chosen over LNG injection due to the prolonged nature of experimental procedures. LNG capsules were constructed in-house of Corning Silastic tubing (Fisher) with dimensions 3cm L, 1.96mm inner diameter, 3.18mm outer diameter. Implants were filled with ∼30mg LNG powder (MilliporeSigma, St. Louis, MO) and sealed with silastic adhesive. Each rat received two implants in the scapular region in a method derived from Ma et al., 2006, intended to mimic the NORPLANT system and induce an anovulatory state (Croxatto, 1993; Nash et al., 1978). Surgical procedures occurred under a 50 mg/kg Ketamine and 5 mg/kg Xylazine anesthesia cocktail (Patterson Veterinary, Greeley, CO). Once anesthetized, 0.03 mg/kg buprenorphine hydrochloride (Patterson Veterinary) was administered via subcutaneous injection. The subcutaneous LNG capsule was inserted into a small (<1cm) skin incision in the scapular region that was sutured and treated with antibiotic ointment. A subgroup of rats (n = 5) received sham surgeries with Ketamine and Xylazine anesthesia, buprenorphine, scapular skin incision and suture, but no capsule implant. Once consciousness was regained rats were returned to the home cage (with cagemates) and allowed ∼5 days to recover prior to any experimental procedures. The efficacy of the LNG implant to induce anovulation was confirmed by a state of persistent diestrus among implanted rats.

#### Light-food conditioning

Light-food appetitive conditioning procedures were used to determine conditioned orienting and conditioned approach behavior over four consecutive sessions. One day prior to conditioning rats were trained in a 30-minute session to retrieve a foodpellet from the foodcup; 30 foodpellets were delivered on a 60s inter-trial interval (ITI) for this session. Each subsequent conditioning session, except for habituation, consisted of 16 CS-US presentations in which illumination of a 10-second light CS was paired with noncontingent delivery of a foodpellet US. Habituation consisted of 8 unpaired CS presentations and 8 paired CS-US presentations, allowing habituation to the light. All conditioning sessions lasted ∼35-minutes with foodpellet delivery occurring on a variable ITI of 120s +/- 60s.

Orienting behavior was video recorded during the conditioning sessions and later scored by a blinded independent observer. Orienting scores were derived over a 15s sampling period: 5s prior to CS onset (preCS), first 5s of CS presentation (CS1), and second 5s of CS presentation (CS2). Scoring occurred every 1.25s and responses were defined as a rearing response where the front legs of the rat were completely removed from the floor (excluding grooming behavior) regardless of the rat’s position within the chamber. Orienting occurs predominantly in CS1 and food cup approach occurs predominantly in CS2 (Hilz et al., 2019a,b; Olshavsky et al., 2014) – basal preCS scores were subtracted from CS1 and CS2 behavior for orienting and foodcup behavior, respectively.

#### Conditioned place preference

CPP procedures similar to those in Hilz et al., 2019a were used with timing adjustments to allow consideration of estrous cycle effects. The procedure took approximately 2 weeks. CPP consisted of four phases: baseline, amphetamine (AMP) and saline conditioning, preference testing / extinction, and reinstatement. The goal for cycling rats during this procedure was to have animals be either P or M/D (i.e., high or low sex steroid hormone state) in each phase (see Fig. 1 for experimental timeline). For example, a P rat would be in P for baseline, AMP (but not saline) conditioning, the first preference test, and reinstatement. The same would be true for an M/D rat. This was accomplished by allowing several days to pass between each procedure while monitoring estrous cycles. HC-implanted rats underwent procedures on a schedule yoked to cycling conspecifics.

**Fig. 1.**
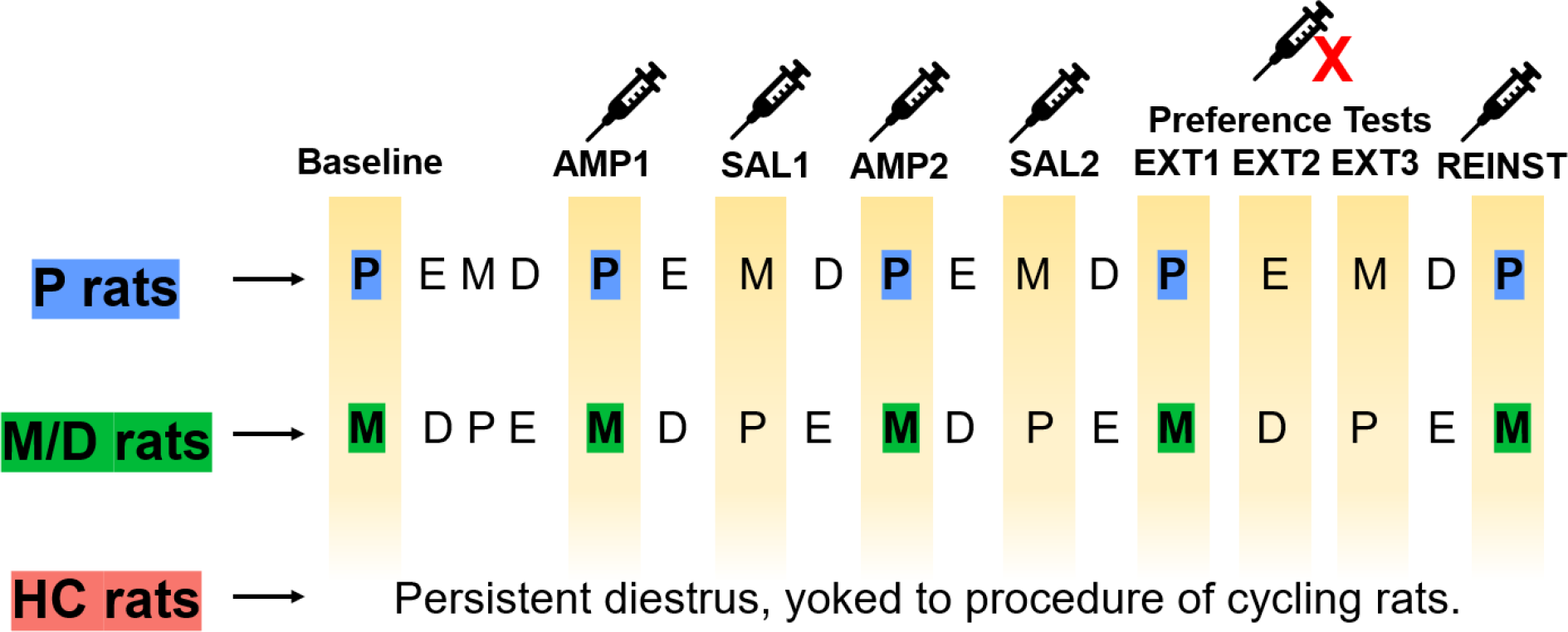
Experimental timeline highlighting the estrous cycle state of intact rats for CPP procedures. Rats cycled freely throughout the procedure and underwent baseline, conditioning (AMP 1 and 2, saline (SAL) 1 and 2), the first day of preference/extinction (EXT) testing, and reinstatement (REINST) in opposing estrous cycle states (P or M/D). Rats with HC implant showed persistent diestrus throughout the procedures and were yoked to the same experimental timeline as cycling rats.

Baseline was a 15-minute session wherein rats were placed into the conditioning chamber on the dark side and allowed to roam freely between the dark and light sides through the open retractable door; baseline determined unconditioned place preference scores. The subsequent conditioning procedures were biased such that amphetamine was paired with the less preferred context. Approximately four days after baseline, once rats had cycled back into P or M/D, conditioning occurred. Conditioning sessions were separated by two days and each consisted of an intraperitoneal (i.p.) injection of either d-amphetamine (AMP; 2.0 mg/kg, Sigma-Aldrich) or 0.9% sterile saline prior to placement within the conditioning chamber with the retractable door closed. The first conditioning session paired AMP with confinement to the less-preferred context for 30 minutes. The second paired saline with confinement to the more-preferred context for 30 minutes. This procedure was repeated such that rats received two AMP and two saline conditioning sessions. Approximately two days after the final conditioning session, once rats had cycled back to P or M/D, preference testing occurred. Rats were either P or M/D for the first preference test. Preference testing occurred over three consecutive days and, as no AMP i.p. injection occurred, acted additionally as extinction training. Preference/extinction tests were 15-minute sessions wherein rats were placed into the conditioning chamber on the dark side and allowed to roam freely between the dark and light sides through the open retractable door. Finally, reinstatement of AMP CPP was assessed via AMP challenge. Reinstatement testing occurred approximately two days after preference / extinction testing, when rats had cycled back to P or M/D. Rats were given AMP i.p. injection prior to placement within the dark chamber and allowed to roam freely through the open retractable door for 15 minutes.

#### Ultrasonic vocalizations

USVs were recorded throughout CPP procedures for a subset of rats (*n* = 28; 5 rats did not have USV recording) and digitized with UltraSoundGate system (Avisoft Bioacoustics) using methods described in Hilz et al., 2021 (in press). USVs were recorded and analyzed using Avisoft-RECORDER and Avisoft-SASlab Pro software (Avisoft Bioacoustics). USVs were analyzed when either saline or AMP were given to the rat (i.e., during the conditioning sessions (AMP or saline pairing) and at reinstatement). Recordings took place over 30 minutes after AMP or saline place conditioning and for 15 minutes after AMP challenge at reinstatement; the CM16 ultrasonic microphones were placed inside the conditioning chamber facing the closed retractable door on the side opposite the confined rat for conditioning and outside the dark chamber for reinstatement. Both flat-type and frequency-modulated (FM) type vocalizations were recorded in accordance with methods from Ahrens et al., (2013); in short, USVs in the 50 kHz range were counted by a blinded independent observer using visual inspection of spectrograms and auditory confirmation. FM-type and flat calls were categorized based on the presence or lack thereof fluctuations in frequency.

#### Histology and immunohistochemistry

Ninety minutes following the CPP reinstatement test rats were anesthetized with 0.3ml pentobarbital and phenytoin (MED-PHARMEX Inc., Pomona, CA) i.p. injection. Blood was collected from the heart and allowed to clot while rats were transcardially perfused with 0.9% saline followed by 4% paraformaldehyde in 0.1 M phosphate buffer. Uterine horn width near the base was also taken as a secondary measure of gonadal hormone production. Blood was centrifuged for 20 minutes after perfusions and the serum was stored at -80 °C. Brains rested for 24 hours at 4°C in 20% sucrose with paraformaldehyde prior to rapid freezing with dry ice and storage at -80°C. Brains were later sliced into 35µm coronal sections (six series) and one series was immediately processed for FOS and tyrosine hydroxylase (FOS+TH).

#### Serum hormone assays

Serum estradiol (E2) was assayed by RIA kit (Beckman Coulter, Webster, TX, #DSL-4800) in a single assay following manufacturer protocols per Yin et al., 2019. The assay range was 2.2 - 750 pg/ul, sensitivity was 2.2 pg/ml, and intra-assay CV was 5.4%. Serum progesterone (P4) was assayed by ELISA kit (Cayman Chemical, Ann Arbor, MI, #582601) in a single assay, as published (Mennigen et al., 2018)..The assay range was 7.8 - 1000 pg/ul, sensitivity was 7.8 pg /ul, and intra-assay CV was 2.86%. Nine rats (P = 4, M/D = 5) were above the standard curve (> 100 pg/ml) for the P4 ELISA.

FOS immunohistochemistry was conducted using procedures identical to Hilz et al., 2019a. In short, primary incubation used rabbit antiserum against FOS (1:2500; Vector Laboratories, Inc., Burlingame, CA) and secondary incubation used biotinylated goat anti-rabbit IgG (1:250; Vector Laboratories). VECTASTAIN ABC standard kit (Vector Laboratories) was used to detect biotinylation and then tissue was stained with 3’3 diaminobenzidine (SIGMA-ALDRICH Co., St. Louis, MO) with nickel. FOS tissue was then co-stained for TH wherein primary incubation used mouse antiserum against TH (1:2500; ImmunoStar, Hudson, WI) and secondary incubation used biotinylated goat anti-mouse IgG (1:250). VECTASTAIN ABC standard kit was used and then tissue was stained with 3’3 diaminobenzidine only.

#### Imagine Acquisition and Analysis

FOS+TH images were acquired under 20x magnification using a Nikon compound light microscope. Levels of the substantia nigra (SN) and the ventral tegmentum area (VTA) were identified using Swanson’s atlas (2004). Images were taken bilaterally at levels 36 and 37 (-5.00mm and -5.25mm from bregma); four samples were taken for SN (two samples at each coordinate) and two samples were taken for VTA (one sample at each coordinate). Images were analyzed by a blinded observer.

### Statistical analyses

#### Light-food conditioning

OR and food cup responses were analyzed using repeated measures factorial ANOVA with the factors Group (i.e., cycling or HC – cycle was not controlled during light-food conditioning) and Training block (i.e., blocks 2-8, 8 CS-US presentations per block). Training block 1 consisted of unpaired CS presentations (analyses not reported). *Post hoc* analyses were conducted using either Tukey HSD for between-subjects’ analyses or Bonferroni corrected *t* tests for repeated measures analyses; effect sizes were calculated using partial eta-squared (η^2^_p_) for significant ANOVAs and Cohen’s *d* (*d* for equal sample sizes and *d_s_* for unequal sample sizes) for significant pair-wise comparisons. These statistical methods were used in this and all subsequent analyses.

#### Conditioned place preference

AMP place preference scores were generated by subtracting time (s) spent in the AMP-associated context at baseline from individual preference/extinction testing sessions. Data were analyzed using separate ANOVAs to assess different aspects of the CPP paradigm: difference in AMP place preference from beginning to end of extinction was analyzed using repeated measures factorial ANOVA with factors Group (i.e., P, M/D, or HC) and Extinction Session (i.e., EXT 1 - 3). Reinstatement of AMP preference is indicated by a return of conditioned responding after extinction training; to analyze this a repeated measures factorial ANOVA was used comparing AMP place preference at the end of extinction compared to at the reinstatement test and included factors Group and Condition (i.e., EXT 3 and REINST).

#### Ultrasonic Vocalizations

USVs were monitored during all CPP experimental procedures but were analyzed only when AMP or saline were given to the rat (i.e., conditioning and REINST). USV response to AMP and saline during conditioning was analyzed using separate repeated measures factorial ANOVA with the factors Group, Type (i.e., flat or FM), and Treatment (i.e., saline 1 and AMP1, or saline 2 and AMP 2). Reinstatement was assessed using an AMP challenge and was analyzed with repeated measures ANOVA with the factors Group and Type. Several rats (*n* = 5) were excluded from analysis due to no USV recording. This same subset of rats’ data was excluded for postmortem physiological measures.

#### TH-FOS immunohistochemistry

Total number of TH+ cells and percentage of TH cells double labelled with FOS in the SN and VTA were analyzed using separate one-way ANOVAs with the factor Group for both SN and VTA.

#### Orienting phenotype in female AMP-response measures

Orienting phenotype was determined using median split on the final acquisition block (i.e., block 8) and both OR and FC behavior was analyzed over conditioning with separate repeated measures factorial ANOVAs with the factors Phenotype (i.e., Orienter or Nonorienter) and Block. The conditioned orienting phenotype was then explored as a potential moderator for the multiple AMP-response measures (i.e., AMP place preference, USV response, and DA response). Analyses were conducted as described above with Phenotype included as a factor.

#### Postmortem physiological levels

Uterine horn thickness, serum estradiol, and serum progesterone concentrations were analyzed using one-way ANOVAs with the factor Group.

#### Regression analyses

Simple linear regressions were used to identify predictors of time spent in the AMP-associated context (normalized with Z-scores) at the end of conditioning (i.e., EXT3) and at reinstatement. A separate set of simple linear regressions were used to identify predictors of both DA activity (i.e., TH+ cells co-labelled with FOS) and availability (i.e., mean TH+ cells).

## Results

### Light-food conditioning

Groups did not show a significant change in conditioned ORs over acquisition blocks 2-8; however, a non-significant trend with a moderate effect size suggested an increase in ORs over Training Blocks (*F*(6,179) = 1.966, *p* = .07, η^2^_p_ = .06) at approximately the same rate between groups indicated by no interaction effect (Fig. 2 Top). Both HC and cycling rats acquired FC responses over acquisition blocks 2-8 at approximately the same rate indicated by a significant main effect of Training Block (*F*(6,186) = 50.02, *p* < .001, η^2^_p_ = 0.62) but no interaction between Group and Training Block (Fig. 2 Bottom).

**Fig. 2.**
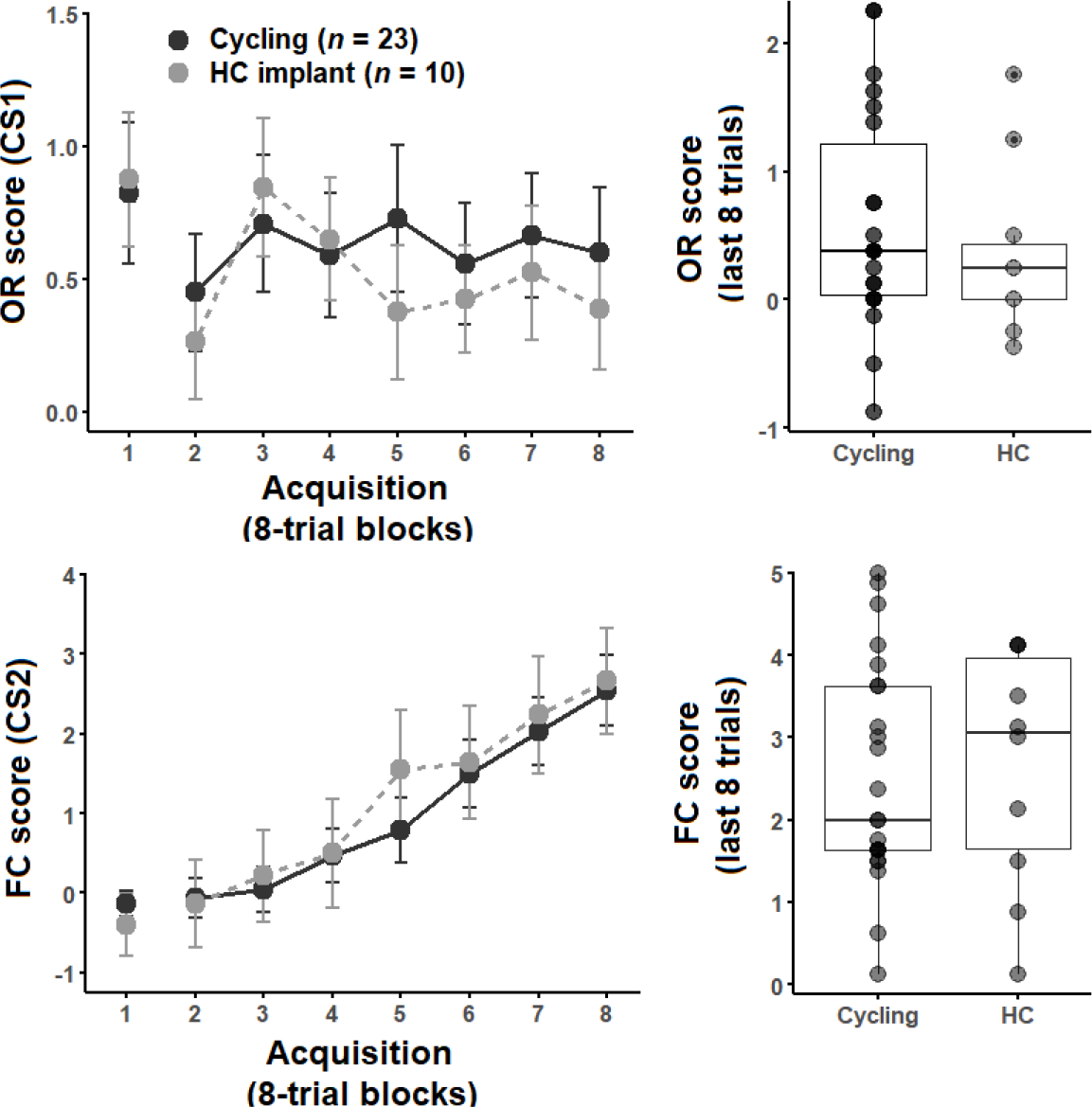
Top Left. Orienting response (OR) score +/- SEM over acquisition. Block 1 is unpaired CS presentations. Groups did not acquire ORs over acquisition. **Top Right.** Boxplot showing spread of OR scores at end of acquisition (block 8). Horizontal line indicates median OR score with upper and lower 50 percentiles. Individual mean scores are represented by points. **Bottom Left.** Foodcup (FC) score +/- SEM over acquisition. Groups acquired FC behavior at approximately the same rate. **Bottom right.** Boxplot showing spread of FC scores at end of acquisition (block 8) with individual scores indicated by points.

### Conditioned place preference

A significant main effect of Group suggested an overall difference in AMP place preference over preference/extinction testing (*F*(2,30) = 3.24, *p* = .05, η^2^_p_ = 0.18) but no effect of extinction session or interaction between session and group were observed (Fig. 3 Left). *Post hoc* analyses with a Bonferroni corrected alpha of *p* < .016 comparing mean time spent in the AMP-associated context over all preference/extinction testing sessions indicated that P rats appeared to spend more time in the AMP-associated context compared to HC (*p* = .04, *d* = 0.81) and M/D (*p* = .03, *d_s_* = 0.75) rats, although these comparisons were not significant at the corrected level (Fig. 3 Right); M/D and HC rats did not significantly differ from each other. No reinstatement of AMP preference was detected when comparing EXT3 to REINST; however, a non-significant interaction between Group and Condition indicated that moderate differences in reinstatement of AMP preference may have occurred depending on Group (*F*(2,30) = 3.12, *p* = .06).

**Fig. 3.**
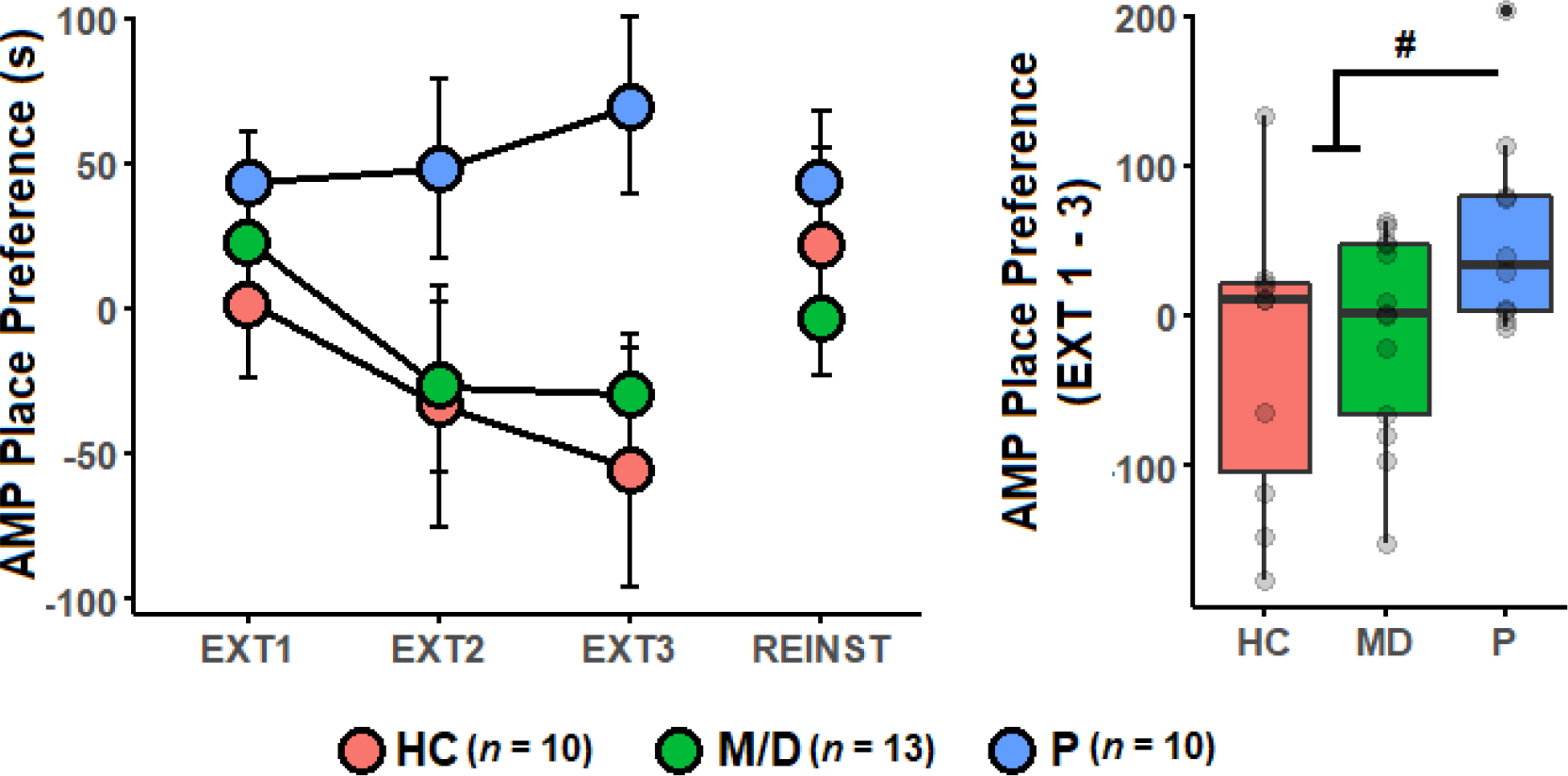
Left. Difference in time spent in the AMP-associated context (seconds (s)) from baseline +/- SEM over preference/extinction testing (i.e., EXT1-3) and at reinstatement (REINST). A significant main effect suggested P rats spent more time in the AMP-associated context over EXT1-3. **Right.** Distribution of mean AMP place preference scores from EXT1-3 with upper and lower quartiles and median (middle line). Individual AMP place preference scores for each rat (points) are included. P rats showed a non-significant trend towards spending more time in the AMP-associated context compared to both M/D and HC rats (# = *p* < .05, nonsignificant after correcting for multiple comparisons).

### Ultrasonic Vocalizations

Significant differences were detected in the number of flat or FM-type USVs emitted during AMP-saline place conditioning (i.e., session 1 and session 2, respectively) indicated by main effects of Group (*F*(2,25) = 3.59, *p* < .05, η^2^_p_ = 0.22; *F*(2,25) = 5.52, *p* = .01, η^2^_p_ = 0.30), Condition (*F*(1,299) = 101.38, *p* < .001, η^2^_p_ = 0.26; *F*(1,299) = 93.63, *p* < .001, η^2^_p_ = 0.24), and Type (*F*(1,299) = 162.02, *p* < .001, η^2^_p_ = 0.36; *F*(1,299) = 177.07, *p* < .001, η^2^_p_ = 0.37; Fig. 4). In addition, significant interactions between Group, Condition, and Type (*F*(2,299) = 8.31, *p* < .001, η^2^_p_ = 0.05; *F*(2,299) = 16.12, *p* < .001, η^2^_p_ = 0.10) indicated differences in USVs emitted between experimental groups depending on call type and conditioning session. *Post hoc* analyses with a Bonferroni corrected alpha of *p* < .008 for between subjects’ interaction effects indicated P rats emitted significantly more FM USVs than HC and M/D rats at AMP 1 (both *p* < .001, *d* = 1.24, *d_s_* = 0.87) and at AMP 2 (both *p* < .001, *d* = 1.07, *d_s_* = 1.46). At reinstatement, only a significant main effect of Type was observed (*F*(1,25) = 38.38, *p* < .001, η^2^_p_ = 0.60) such that more FM calls were emitted compared to flats; no significant differences were observed in USVs between groups at reinstatement.

**Fig. 4.**
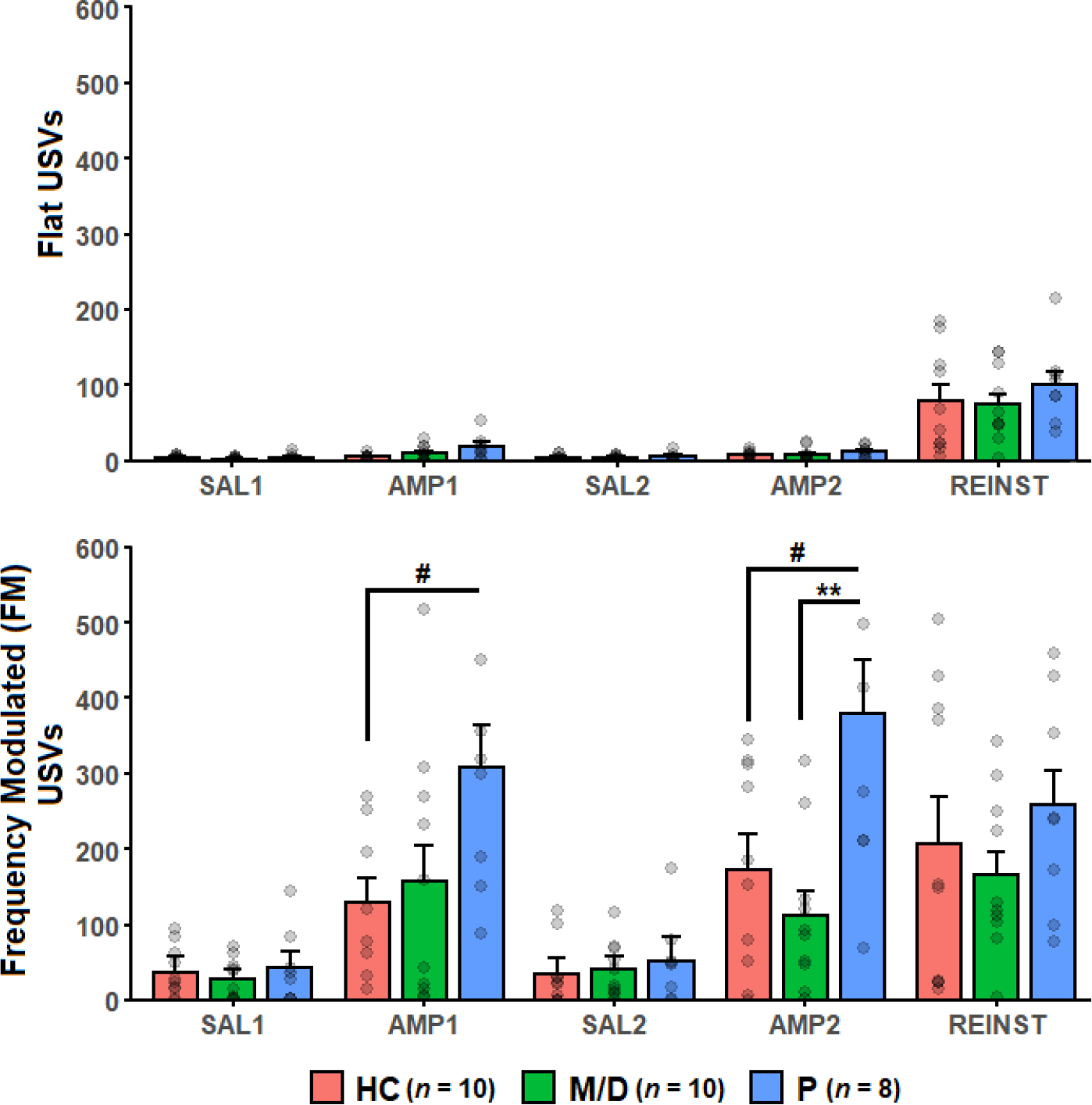
Mean flat (top) or frequency modulated (FM) type (bottom) USVs +/- SEM. Individual USV scores indicated by gray points. **Top.** No differences were observed between groups in flat-type USVs. More flats were observed at REINST than during conditioning. **Bottom.** All groups emitted significantly more FM USVs in response to AMP than to saline (SAL). P rats emitted significantly more FM-type USVs compared to M/D and HC rats at both AMP1 and AMP2.

#### TH-FOS immunohistochemistry

The total number of TH+ cells in the SN differed depending on Group (*F*(2,233) = 4.42, *p* < .05, η^2^_p_ = 0.04); *post hoc* analyses using Tukey’s HSD indicated that P rats had significantly more overall TH+ cells compared to HC (*p* < .05, *d* = 0.46) but not M/D (*p* = .06, *d_s_* = 0.36) rats (Fig. 5 Left). The percentage of active TH+ cells also differed between groups in the SN (*F*(2,233) = 12.69, *p* < .001, η^2^_p_ = 0.10) with P rats having a higher percentage of active TH+ cells compared to both M/D (*p* < .001, *d_s_* = 0.69) and HC (*p* < .001, *d* = 0.65) rats (Fig. 5 Right). In the VTA neither total number of TH+ cells or percentage of active TH+ cells differed between groups (*F*(2,117) = 0.69; *F*(2,117) = 0.84).

**Fig 5.**
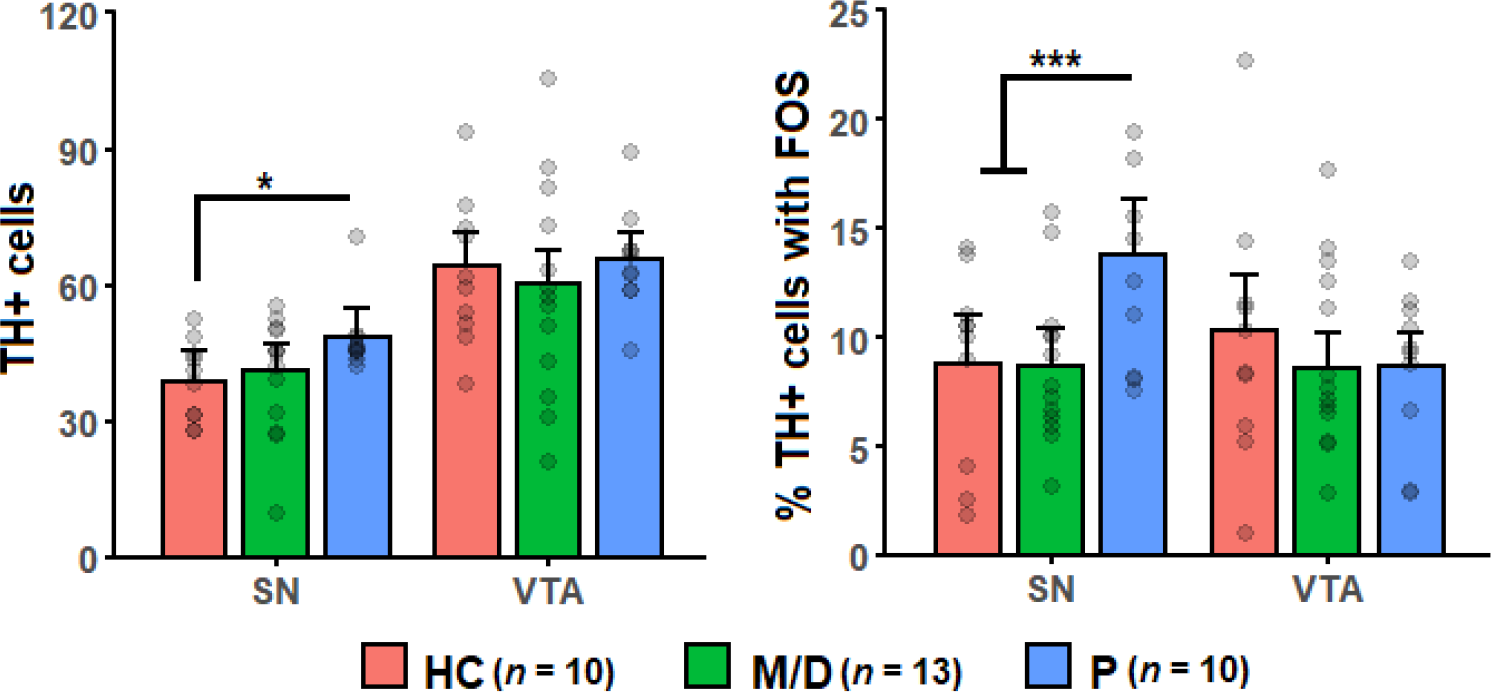
Mean TH+ cells (left) and mean percent TH+FOS cells (right) +/- SEM. **Left.** HC rats had significantly fewer TH+ cells in SN compared to P rats. **Right.** P rats had a significantly higher percent of TH+ cells with FOS in SN compared to M/D and HC rats.

#### Orienting phenotype in female AMP-response measures

The median of OR scores at block 8 was 0.375, consistent with both published and unpublished work from our lab (Olshavsky et al., 2014). Rats with an OR score above the median were labeled ‘Orienters’, while those below were labeled ‘Nonorienters’. A significant main effect of Phenotype (*F*(6,186) = 4.40, *p* < .001, η^2^_p_ = 0.13) and of Block (*F*(6,186) = 2.19, *p* < .05, η^2^_p_ = 0.07), along with a significant interaction between Phenotype and Block (*F*(1,31) = 21.58, *p* < .001, η^2^_p_ = 0.43), indicated acquisition of ORs over blocks 2-8 significantly differed between Orienters and Nonorienters (Fig. 6 Left); *post hoc* tests with an adjusted alpha of *p* < .025 conducted on final acquisition block 8 indicated Orienters expressed significantly more ORs at the end of conditioning than Nonorienters (*p* < .001, *d_s_* = 2.00; Fig. 6 Right). Orienters and Nonorienters acquired foodcup behavior at approximately the same rate, indicated by a main effect of Block (*F*(6,186) = 49.20, *p* < .001, η^2^_p_ = 0.61) but no effect of Phenotype or interaction between Phenotype and Block (data not shown). These designations were used to determine if the orienting phenotype moderated female AMP-response measures by including Phenotype as potential moderator of female AMP responsivity (i.e., AMP place preference, USV response, and DA response). Generally, orienting phenotype did not significantly affect female AMP response measures (see Table 1). Phenotype did interact with Group for percentage of TH+ cells co-labelled with FOS (i.e., active DA) in the SN (*F*(2,230) = 10.65, *p* < .001, η^2^_p_ = 0.08). In the SN, the percentage of TH+ cells co-labelled with FOS was significantly higher among P rats with an Orienter phenotype compared to all other groups (*p* < .001 all; supplemental figure 1).

**Fig. 6.**
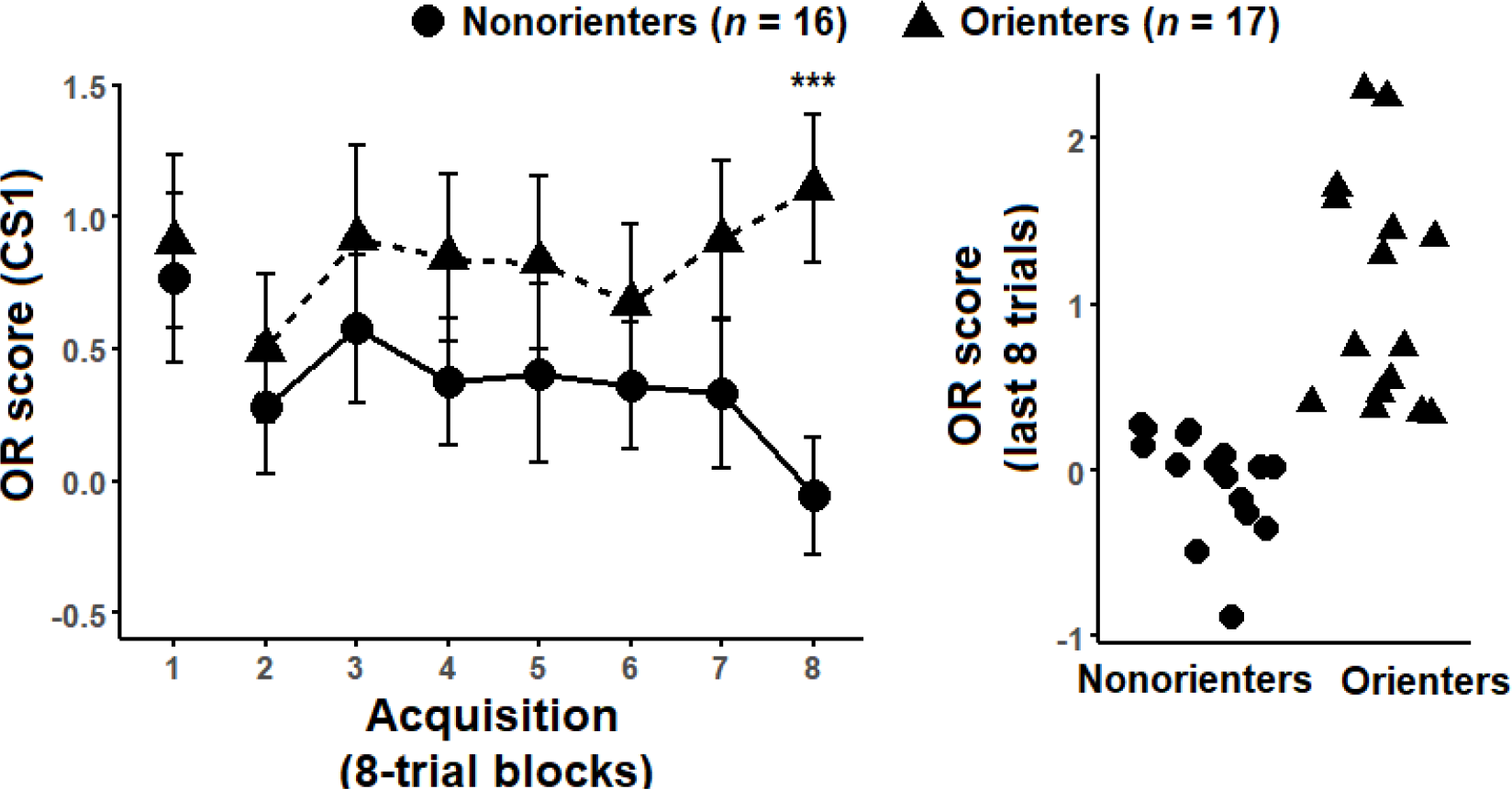
Left. OR score +/- SEM. Orienters significantly acquired OR scores over acquisition blocks and had a significantly higher OR score at the end of acquisition compared to Nonorienters. **Right.** Scatterplot showing individual OR scores of Orienters and Nonorienters.

**Table 1.** Conditioned orienting in female AMP, USV, and DA measures.

| *Amphetamine (AMP) place preference* |  |  | *Ultrasonic Vocalizations (USVs)* |  |  |
| --- | --- | --- | --- | --- | --- |
| *Initial AMP place preference (EXT1)* |  |  | *Conditioning 1 (SAL1 and AMP1)* |  |  |
| Phenotype | $F(1,27) = 1.17$ | $> .1$ | Phenotype | $F(1,22) = 0$ | $> .1$ |
| Group | $F(2,27) = 0.72$ | $> .1$ | Group | $F(2,22) = 3.04$ | $0.07\#$ |
| (Phenotype*Group) | $F(2,27) = 0.62$ | $> .1$ | (Phenotype*Group*Condition*Type) | $F(2,290) = 2.07$ | $> .1$ |
| *Preference/extinction testing (EXT1-3)* |  |  | *Conditioning 2 (SAL2 and AMP2)* |  |  |
| Phenotype | $F(1,27) = 0.6$ | $> .1$ | Phenotype | $F(1,22) = 0.06$ | $> .1$ |
| Group | $F(2,27) = 3.19$ | $= .06$ | Group | $F(2,22) = 5.20$ | $= .01^{**}$ |
| (Phenotype*Group) | $F(2,27) = 0.96$ | $> .1$ | (Phenotype*Group*Condition*Type) | $F(2,290) = 0.64$ | $> .1$ |
| *Reinstatement (EXT3 v. REINST)* |  |  | *Reinstatement* |  |  |
| Phenotype | $F(1,27) = 0.006$ | $> .1$ | Phenotype | $F(1,22) = 0.20$ | $> .1$ |
| Group | $F(2,27) = 3.18$ | $= .06$ | Group | $F(2,22) = 0.70$ | $> .1$ |
| (Phenotype*Group) | $F(2,27) = 1.73$ | $> .1$ | (Phenotype*Group*Type) | $F(2,22) = 0.41$ | $> .1$ |
| *TH and FOS measures* |  |  |  |  |  |
| *Substantia Nigra (SN)* |  |  | *Ventral Tegmental Area (VTA)* |  |  |
| *Percent TH+ cells with FOS* |  |  | *Percent TH+ cells with FOS* |  |  |
| Phenotype | $F(1,230) = 5.24$ | $< .05^*$ | Phenotype | $F(1,114) = 0.95$ | $> .1$ |
| Group | $F(2,230) = 13.97$ | $< .001^{***}$ | Group | $F(2,114) = 0.86$ | $> .1$ |
| (Phenotype*Group) | $F(2,230) = 10.65$ | $< .001^{***}$ | (Phenotype*Group) | $F(2,114) = 5.82$ | $= .08$ |
| *Mean TH+ cells* |  |  | *Mean TH+ cells* |  |  |
| Phenotype | $F(1,230) = 1.32$ | $> .1$ | Phenotype | $F(1,114) = 0.68$ | $> .1$ |
| Group | $F(2,230) = 4.40$ | $= .01^{**}$ | Group | $F(2,114) = 0.69$ | $> .1$ |
| (Phenotype*Group) | $F(1,230) = 0.38$ | $> .1$ | (Phenotype*Group) | $F(2,114) = 0.62$ | $> .1$ |

*Note.* Results of repeated measures ANOVA considering conditioned orienting phenotype as a factor in female AMP place preference, USV response to AMP and saline (SAL), and DA measures.

### Postmortem physiological measures

Serum estradiol (E2) concentrations did not significantly differ between groups (*F*(2,25) = 2.30; Fig. 7 Left). Serum progesterone (P4) concentrations significantly differed between groups (*F*(2,25) = 13.78, *p* < .001, η^2^_p_ = .52) with *post hoc* Tukey HSD indicating HC rats had significantly lower levels of serum P4 compared to both P (*p* < .001, *d* = 2.16) and M/D (*p* < .001, *d_s_* = 2.23) rats (Fig. 7 Middle). Uterine horn width significantly differed between groups (*F*(2,25) = 35.99, *p* < .001, η^2^_p_ = 0.74) with P rats having significantly thicker uterine horn widths compared to M/D (*p* < .001, *d_s_* = 3.02) and HC (*p* < .001, *d_s_* = 3.41) rats (Fig. 7 Right).

**Fig. 7.**
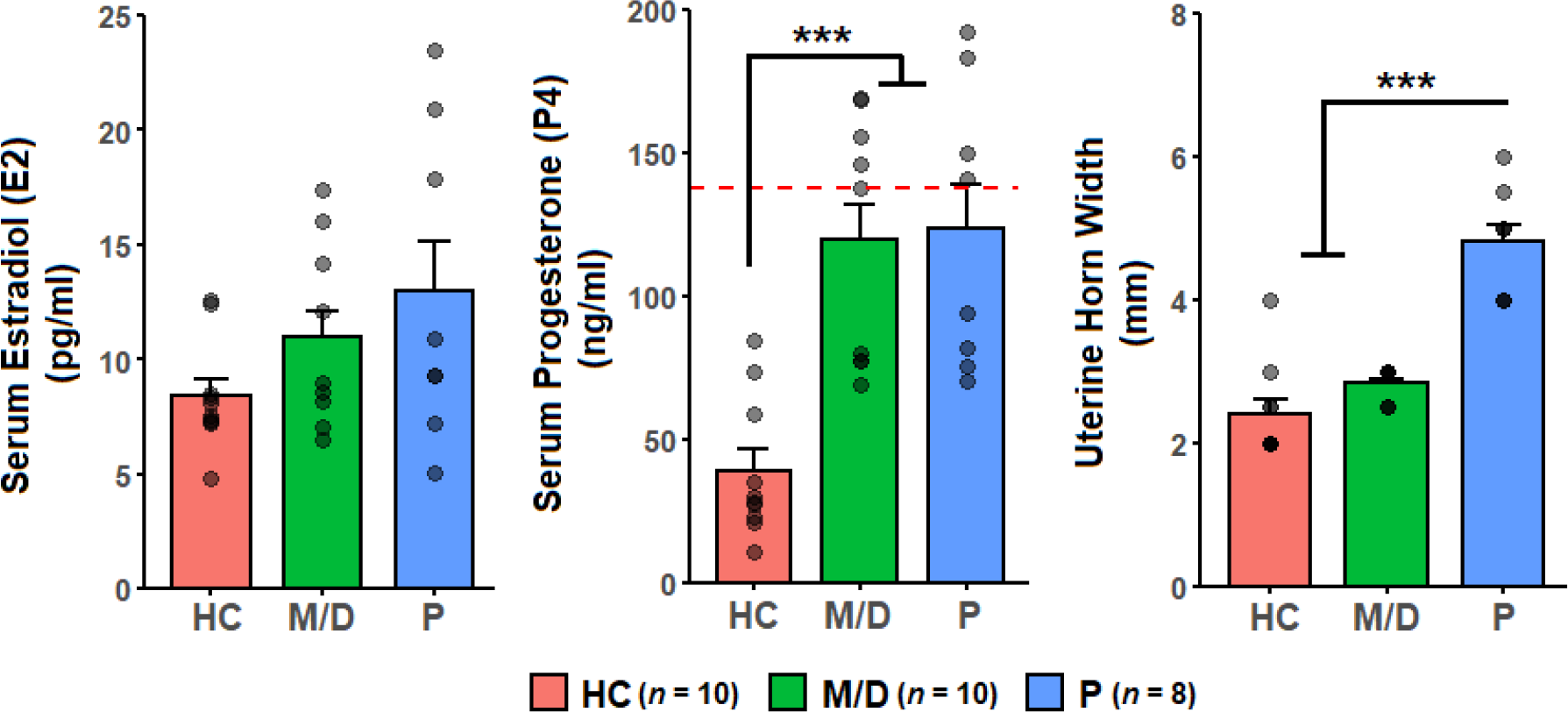
Postmortem physiological measures. Individual data points shown in gray. **Left.** Mean serum estradiol (E2) +/- SEM. Serum E2 did not significantly differ between groups. **Middle.** Mean serum progesterone (P4) +/- SEM. HC rats had lower serum P4 compared to both M/D and HC rats (*p* < .001). Points at or above red dashed line were above standard curve. All data was included for calculating mean and SEM. **Right.** Uterine horn width, an index of prior exposure to estrogens, is shown +/- SEM). Both HC and M/D rats had thinner uteri horn widths compared to P rats (*p* < .001).

### Regression analyses

Potential predictors of time spent in the AMP-associated context at both the end of extinction (i.e., EXT3) and at reinstatement are detailed in supplementary Table 1. Several significant relationships were observed. Percent of DA activity in the SN (R^2^ = 0.15, β = 0.08 +/- 0.03, *p* < .05; Fig. 8A), serum E2 concentration (R^2^ = 0.17, β = 0.09 +/- 0.04, *p* < .05; Fig. 8B), and uterine horn width (R^2^ = 0.20, β =0.38 +/- 0.15, *p* < .05; Fig. 8C) all significantly predicted time spent in the AMP-associated context at EXT3. Additionally, a trend was observed wherein FM USVs emitted during the first AMP conditioning session (i.e., AMP1) also non-significantly predicted time spent in the AMP-associated context at EXT3 (R^2^ = 0.13, *p* = .06, data not shown).

**Fig. 8.**
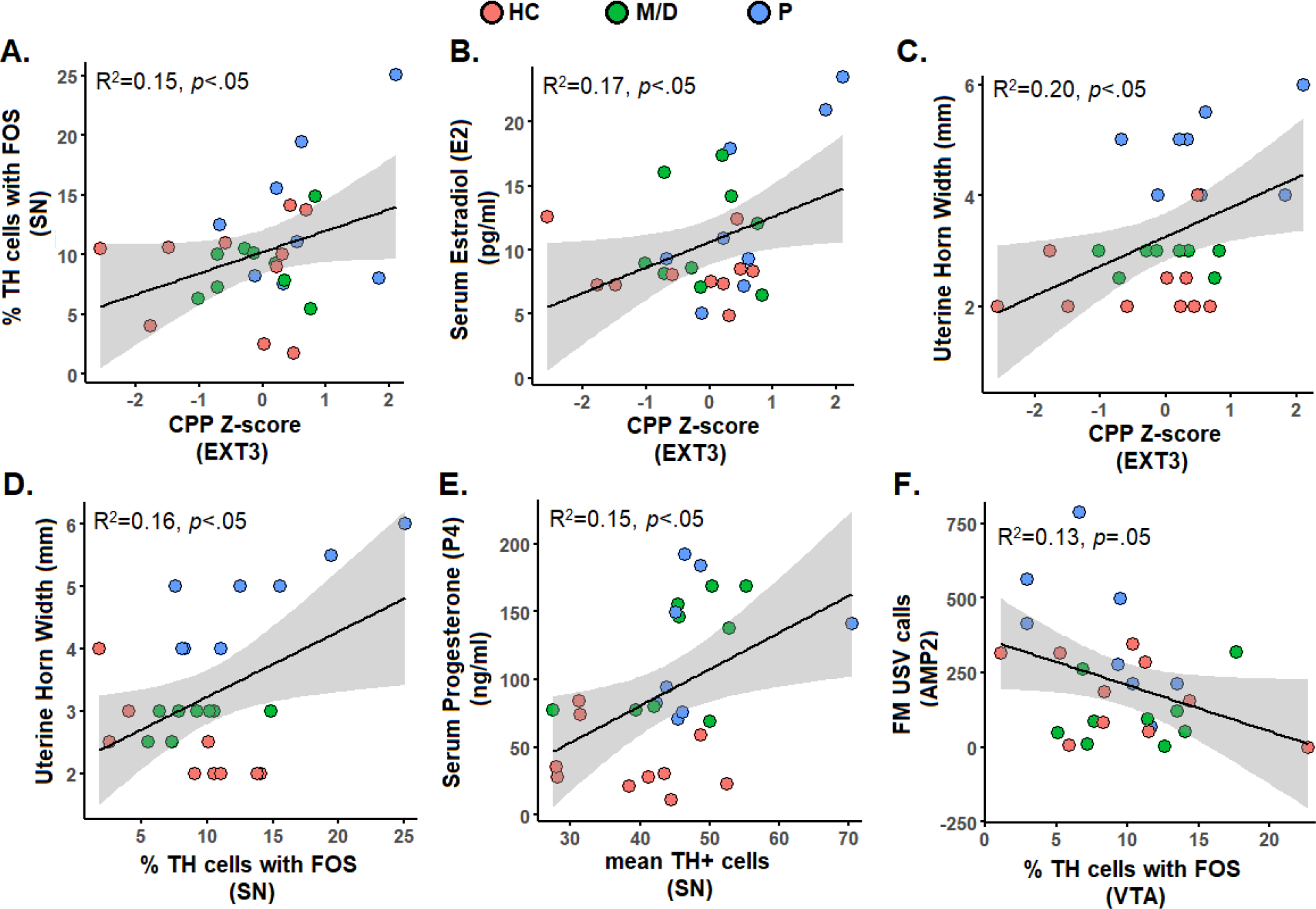
Simple linear regressions showing predictors of AMP place preference Z-scores (top row) and DA measures (bottom row). Individual points for experimental groups are shown along with R^2^ and alpha (*p*). **Top Row.** Rats that spent more time in the AMP-associated context at EXT3 also showed a higher percentage of active DA cells (i.e., TH+ cells co-labelled with FOS) (A), higher serum concentrations of estradiol (i.e., E2) (B), and wider uterine horns (C) after reinstatement. **Bottom Row.** In SN, higher percentages of active DA cells were associated with wider uterine horns (D), and more DA cells (i.e., TH+ cells) were associated with increased serum progesterone (P4) (E). In VTA, higher percentages of active DA cells were associated with fewer frequency-modulated (FM) USVs (F).

These relationships were not observed for reinstatement of AMP place preference, where only the percent of DA activity in SN was a notable non-significant predictor (R^2^ = 0.09, *p* = 0.08). Predictors of DA activity and/or availability were also considered in supplementary Table 2. In SN, the percentage of active DA cells was predicted by uterine horn width (R^2^ = 0.16, β = 1.67 +/- 0.74, *p* < .05; Fig. 8D) and DA availability was predicted by serum P4 concentration (R^2^ = 0.15, β = 0.08 +/- 0.04, *p* < .05; Fig. 8E). In VTA, percent of active DA was predicted by FM USVs emitted during the second AMP conditioning session (i.e., AMP2) (R^2^ = 0.13, β = -.008 +/- 0.004, *p* = .05; Fig. 8F).

## Discussion

The results of these experiments suggest that modulating the estrous cycle of female rats with HCs has important implications for subsequent reward-learning and drug-responsive behavior, and that HCs may be a valuable tool for attenuating female drug preference. HC rats spent less time in the AMP-associated context over preference/extinction testing, emitted fewer USVs in response to AMP place conditioning, and had both a lower number of DA cells and a lower percentage of active DA cells in the SN compared to P rats. For many of these measures HC rats had similar scores as M/D rats, indicating that the effect of HCs on circulating sex steroid hormones behaviorally mimics a low endogenous hormonal state. HC rats had both lower P4 serum concentrations than cycling rats and a similar uterine horn width (a common measure of estradiol responsivity around the estrous cycle; Spornitz et al., 1994) as M/D rats at the time of sacrifice. These physiological measures predicted AMP place preference and DA immunoreactivity after AMP challenge.

### Conditioned orienting in female rats

Conditioned orienting scores did not differ between cycling and HC-implanted rats, indicating that ORs are not influenced by sex steroid hormone availability. This is supported by a recent finding from Hilz et al., 2021 (in press) showing that OVX and/or estradiol replacement did not affect female conditioned orienting; however, sex differences were observed in conditioned orienting such that female rats expressed more ORs than males regardless of estradiol replacement. This result may seem counterintuitive, as conditioned orienting is thought to rely on functional activity of the DA system and the DA system is strongly modulated by ovarian sex steroid hormones. The lack of an effect of sex steroid hormone availability on expression of conditioned ORs may indicate that conditioned orienting is a phenotype organized early in life (Morris et al., 2004). While conditioned orienting did not differ between groups, there were some measures of AMP responsivity that were modulated by the conditioned orienting phenotype. Specifically, the percentage of active DA cells in the SN was significantly higher among P rats with an Orienting phenotype compared to all other groups. Conditioned orienting has also been shown to predict AMP place preference in female rats (Hilz et al., 2019a) and relies on functional activity in the midbrain DA system (Lee et al., 2011; 2005; Han et al., 1997; Gallagher et al., 1990). This experiment did not replicate the finding that the orienting phenotype predicts resistance to extinction of AMP place preference; as this experiment was not designed to detect differences in AMP response measures based on the orienting phenotype, this result may be a product of underpowered and uneven analyses. However, it does provide evidence that DA cell activity is higher among Orienters after AMP challenge and suggests that the orienting phenotype interacts with sex steroid hormones to modulate DA cell activity.

#### The estrous cycle modulates female AMP response measures

While no differences were observed in reinstatement of AMP place preference, there was an overall effect of the estrous cycle on AMP place preference during the extinction test sessions. Rats that were conditioned in P spent more time in the AMP-associated context over preference/extinction testing compared to HC and M/D rats, although these comparisons were non-significant trends after correction for multiple comparisons. In general, this supports previous literature wherein estradiol was shown to enhance drug CPP in OVX models (Mirbaha et al., 2009; Silverman and Koenig, 2007; Russo et al., 2003a,b) and is the first of its kind to replicate sex steroid effects on drug CPP using the estrous cycle. The results suggest that elevated sex steroid hormones associated with the P stage of the estrous cycle promote higher overall AMP place preference and resistance to extinction of this preference compared to rats with artificial abolition of estrous cycling (i.e., HC rats) and rats in low sex steroid hormonal states (i.e., M/D rats). Although the comparison was non-significant, a moderately large effect size gives it credence. Additionally, AMP place preference at the end of extinction was predicted by serum estradiol concentration and uterine horn width. Moreover, groups did significantly differ on other measures of AMP responsivity including FM USVs during AMP place conditioning and DA cell activity after the AMP challenge at reinstatement. For both, P rats exhibited a higher AMP response.

USVs emitted in the 50 kHz range are thought to represent a positive affective state and are emitted by rats in response to AMP (Burgdorf et al., 2016; Wright et al., 2010). Few papers have examined the effects of female sex steroid hormones on USV response to AMP; generally, it appears that intact female rats show robust USV response to AMP, but OVX reduces this responsivity and estradiol replacement fails to rescue it (Hilz et al., 2021, in press). Here, clear effects of estrous cycle modulation on AMP USV responsivity were observed wherein P rats emitted significantly more USVs than M/D and HC rats during AMP place conditioning. It is difficult to determine the precise mechanism via which the estrous cycle modulates AMP USV responsivity. Hilz et al. 2021 (in press) previously showed that estradiol treatment did not enhance AMP USV response; there is evidence that progesterone may underlie this relationship, as progesterone treatment of estradiol primed animals enhances DA response to AMP (Becker and Rudick, 1999) and interacts with DA to mediate sexual receptivity in methamphetamine treated rats (Holder et al., 2015). HC rats showed both lower AMP USV response and reduced serum progesterone; however, while M/D rats also had a reduced AMP USV response, serum progesterone levels did not significantly differ from P rats. Moreover, none of the postmortem physiological measures were good predictors of AMP USV responsivity (data not reported), but AMP USV during initial place conditioning was a modest non-significant predictor of AMP place preference at the end of extinction. Interestingly, FM USVs at the second AMP place conditioning session had an inverse predictive relationship with the percentage of active DA cells after AMP challenge and indicated that as DA activity increased the number of FM USVs decreased. This relationship was not reflected in other AMP conditioning or reinstatement USV measures but could indicate that the USV response to AMP is not a reflection of increased incentive-motivation.

Although rats did not differ in AMP place preference at reinstatement, patterns of DA cell immunoreactivity after the reinstatement AMP challenge significantly differed between groups. HC rats had a reduced number of total DA cells in the substantia nigra (SN) compared to P rats, and both M/D and HC rats had a lower percentage of active DA cells in the SN compared to P rats. The reduced DA cell availability in the SN could suggest that HC-implants may have long-term impacts on DA availability. There is evidence that long-term deprivation of female sex steroid hormones reduces various measures of DA (Kipp et al., 2006; Dluzen, 2000; Leranth et al., 2000; Bossé et al., 1997; Morissette and Di Paolo, 1993). While a reduction was seen in DA cell availability among HC rats – the size of the effect was small-to-modest and not observed between HC and M/D rats. As M/D rats also undergo estrous cycling and should have similar sex steroid hormone availability as P rats (albeit at different, unobserved timepoints), this result may be a facet of an outlier among P rats. However, credence is given to the result by the predictive relationship between serum progesterone level and SN DA cell availability, although with multiple rats showing unusually high serum P4 levels above the standard curve this interpretation should be approached with caution. As LNG is a progestin HC and binds with high affinity to progesterone receptors (Lemus et al., 1992), what may be observed here is an effect of progesterone availability on long-term DA cell availability.

The differences observed in DA cell activity after AMP challenge were more robust and suggest that DA response to the AMP challenge at reinstatement was increased by elevated sex steroid hormones associated with the P stage of the estrous cycle. This is particularly clear when observing the predictive relationship between postmortem physiological measures and DA cell immunoreactivity: DA cell activity was well-predicted by uterine horn width, a more static measure of estradiol responsivity around the estrous cycle. These differences were seen in the SN but not the VTA; both regions are important for reward-seeking and reward-responsive behavior and have equal distribution of sex steroid hormone receptors (Willing and Wagner, 2016; Yoest et al., 2014; Wise, 2009; Gréco et al., 2001). It is unclear why SN was differentially responsive over VTA, or if that difference is particularly meaningful as SN and VTA are considered to have both morphological and functional overlap (Wise 2009; Quinlan et al., 2004; Schultz, 1998). Regardless, activity of DA cells in the SN was a modest non-significant predictor of reinstatement of AMP CPP and a significant predictor of AMP place preference at the end of extinction.

#### Sex steroid measures

Serum estradiol and progesterone concentrations were differentially impacted by the HC implant: serum estradiol concentrations were similar among groups, but rats with an HC implant had lower serum progesterone concentrations compared to both M/D and P rats. Levonorgestrel, the HC used in this experiment, binds with high affinity to progesterone and androgen, but not estrogen, receptors (Lemus et al., 1992). The result that HC rats had lower serum progesterone concentrations compared to cycling rats was not surprising; however, proestrus is typically associated with higher serum estradiol and progesterone concentrations compared to metestrus and early diestrus, and this result was not observed (although several M/D and P rats measured P4 levels above of the standard curve and this result may be driven by that). The timing of measurement is imperative to detect differences in sex steroid hormone availability across the estrous cycle: peaks in estradiol and progesterone only co-occur for a short period of time (with estradiol peak preceding progesterone and then rapidly declining to floor levels); typically, both peaks occur in the light phase of the dark-light cycle (Smith et al., 1975; Butcher et al., 1974). Therefore, it is plausible that peak estradiol and/or progesterone associated with proestrus was missed in this study. The finding that uterine horn width was thickest in proestrus rats indicates prior exposure to high estradiol based on its uterotrophic effect (Spornitz et al., 1994; Ford, 1982). Here, all procedures, including serum collection, occurred approximately 1-3 hours after dark onset. The differences seen in behavior and DA activity were both directly and indirectly indicative of a sex steroid hormone involvement; serum concentrations and physiological measures had predictive relationships with the CPP and DA measures, and estrous/HC groups were differential in responding in a manner that corresponded to inferred gonadal hormone state. This suggests that differential effects of estradiol and/or progesterone on behavioral and DA responsivity to AMP may persist despite a lack of observed serum concentration differences between cycling rats, perhaps after P rats have returned to basal serum concentrations. There is some support for persistent responsivity of DA after estradiol treatment lasting up to 12 hours (Becker and Rudick, 1999). The various relationships between sex steroid measures, behavior, and DAergic cell response provides strong evidence for a role of female gonadal hormonal states in drug preference.

In summary, the current study demonstrates a role of sex steroid hormones and their modulation by hormonal contraceptives in female drug preference and related response measures and indicates that endogenous sex steroids may increase the likelihood of substance abuse perhaps via an interaction of enhanced incentive-motivation (DA response) for substances and increased hedonic pleasure (USV response) in response to substances; HCs could aid treatment of problematic drug use by reducing these measures. It is apparent that the hormonal state of the female learner during reward-learning is important to consider when identifying how female sex steroid hormones will contribute to behavioral outcomes.

## Supplemental Graphs and Tables

**Supplemental figure 1.**
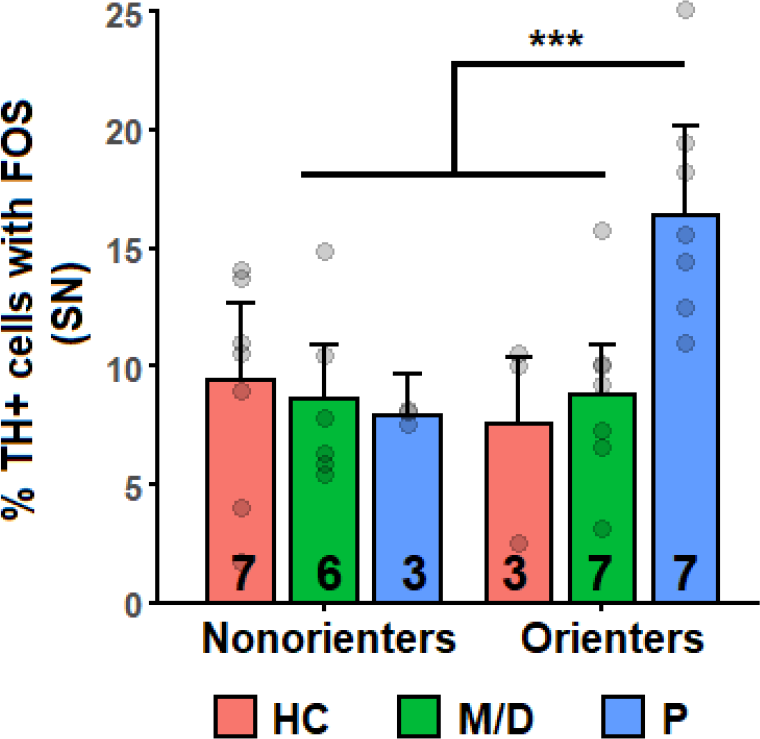
Mean percent of TH+ cells in SN +/- SEM. Gray points = individual scores, numbers on bars = *n* size. P rats with an Orienter phenotype had a significantly higher percentage of TH+ cells in SN compared to all other groups.

**Supplemental Table 1.** Predictors of AMP place preference.

|  | R<sup>2</sup> | β | p |
| --- | --- | --- | --- |
| *Time spent in AMP-associated context (EXT3)* |  |  |  |
| Percent TH+ cells with FOS |  |  |  |
| SN | 0.15 | 0.08 ± 0.03 | < .05* |
| VTA | 0.05 | --- | > .1 |
| Mean TH+ cells |  |  |  |
| SN | 0.01 | --- | > .1 |
| VTA | 0.008 | --- | > .1 |
| OR score | 0.001 | --- | > .1 |
| Physiological measures |  |  |  |
| E2 | 0.17 | 0.09 ± 0.04 | < .05* |
| P4 | 0 | --- | > .1 |
| Uterine Thickness | 0.2 | 0.38 ± 0.15 | < .05* |
| Ultrasonic Vocalizations |  |  |  |
| AMP1 | 0.13 | 0.002 ± 0.001 | = .06# |
| AMP2 | 0.06 | --- | > .1 |
| REINST | 0.003 | --- | > .1 |
| *Time spent in AMP-associated context (REINST)* |  |  |  |
| Percent TH+ cells with FOS |  |  |  |
| SN | 0.09 | 0.06 ± 0.03 | = .08 # |
| VTA | 0.0004 | --- | > .1 |
| Mean TH+ cells |  |  |  |
| SN | 0.0003 | --- | > .1 |
| VTA | 0.0003 | --- | > .1 |
| OR score | 0.01 | --- | > .1 |
| Physiological measures |  |  |  |
| E2 | 0.02 | --- | > .1 |
| P4 | 0.03 | --- | > .1 |
| Uterine Thickness | 0.006 | --- | > .1 |
| Ultrasonic Vocalizations |  |  |  |
| AMP1 | 0.003 | --- | > .1 |
| AMP2 | 0.02 | --- | > .1 |
| REINST | 0.09 | --- | = .1 |

*Note.* Relationships between time spent in the AMP-associated context (difference from baseline) at EXT3 and REINST and related response/physiological measures using simple linear regression.

**Supplemental Table 2.** Predictors of TH and FOS Immunohistochemistry.

|  | R<sup>2</sup> | β | p |  | R<sup>2</sup> | β | p |
| --- | --- | --- | --- | --- | --- | --- | --- |
| *Percent TH+ cells with FOS (SN)* |  |  |  | *Mean TH+ cells (SN)* |  |  |  |
| OR score | 0.03 | --- | > .1 | OR score | 0.01 | --- | > .01 |
| Physiological measures |  |  |  | Physiological measures |  |  |  |
| E2 | 0.03 | --- | > .1 | E2 | 0.03 | --- | > .1 |
| P4 | 0.05 | --- | > .1 | P4 | 0.15 | 0.08 ± 0.04 | < .05* |
| Uterine Thickness | 0.16 | 1.67 ± 0.74 | < .05* | Uterine Thickness | 0.02 | --- | > .1 |
| Ultrasonic Vocalizations |  |  |  | Ultrasonic Vocalizations |  |  |  |
| AMP1 | 0.0004 | --- | > .1 | AMP1 | 0.02 | --- | > .1 |
| AMP2 | 0.01 | --- | > .1 | AMP2 | 0.01 | --- | > .1 |
| REINST | 0.03 | --- | > .1 | REINST | 0.0009 | --- | > .1 |
| *Percent TH+ cells with FOS (VTA)* |  |  |  | *Mean TH+ cells (VTA)* |  |  |  |
| OR score | 0 | --- | > .1 | OR score | 0.0008 | --- | > .1 |
| Physiological measures |  |  |  | Physiological measures |  |  |  |
| E2 | 0.005 | --- | > .1 | E2 | 0.05 | --- | > .1 |
| P4 | 0.002 | --- | > .1 | P4 | 0.02 | --- | > .1 |
| Uterine Thickness | 0.01 | --- | > .1 | Uterine Thickness | 0.005 | --- | > .1 |
| Ultrasonic Vocalizations |  |  |  | Ultrasonic Vocalizations |  |  |  |
| AMP1 | 0.07 | --- | > .1 | AMP1 | 0.01 | --- | > .1 |
| AMP2 | 0.13 | -.008 ± 0.004 | = .05* | AMP2 | 0.003 | --- | > .1 |
| REINST | 0 | --- | > .1 | REINST | 0.006 | --- | > .1 |

*Note.* Relationships between DA cell activity (percent TH cells with FOS) and DA cell availability (mean TH+ cells) and related response/physiological measures using simple linear regression.

